# Recommendations for application of the functional evidence PS3/BS3 criterion using the ACMG/AMP sequence variant interpretation framework

**DOI:** 10.1101/709428

**Authors:** Sarah E. Brnich, Ahmad N. Abou Tayoun, Fergus J. Couch, Garry R. Cutting, Marc S. Greenblatt, Christopher D. Heinen, Dona M. Kanavy, Xi Luo, Shannon M. McNulty, Lea M. Starita, Sean V. Tavtigian, Matt W. Wright, Steven M. Harrison, Leslie G. Biesecker, Jonathan S. Berg, On behalf of the Clinical Genome Resource Sequence Variant Interpretation Working Group

**Author notes:** Contact: Jonathan S. Berg, MD, PhD, 120 Mason Farm Rd., Chapel Hill NC, 27599-7264.

## Abstract

**Background:** The American College of Medical Genetics and Genomics (ACMG)/Association for Molecular Pathology (AMP) clinical variant interpretation guidelines established criteria (PS3/BS3) for functional assays that specified a “strong” level of evidence. However, they did not provide detailed guidance on how functional evidence should be evaluated, and differences in the application of the PS3/BS3 codes is a contributor to variant interpretation discordance between laboratories. This recommendation seeks to provide a more structured approach to the assessment of functional assays for variant interpretation and guidance on the use of various levels of strength based on assay validation.

**Methods:** The Clinical Genome Resource (ClinGen) Sequence Variant Interpretation (SVI) Working Group used curated functional evidence from ClinGen Variant Curation Expert Panel-developed rule specifications and expert opinions to refine the PS3/BS3 criteria over multiple in-person and virtual meetings. We estimated odds of pathogenicity for assays using various numbers of variant controls to determine the minimum controls required to reach moderate level evidence. Feedback from the ClinGen Steering Committee and outside experts were incorporated into the recommendations at multiple stages of development.

**Results:** The SVI Working Group developed recommendations for evaluators regarding the assessment of the clinical validity of functional data and a four-step provisional framework to determine the appropriate strength of evidence that can be applied in clinical variant interpretation. These steps are: 1. Define the disease mechanism; 2. Evaluate applicability of general classes of assays used in the field; 3. Evaluate validity of specific instances of assays; 4. Apply evidence to individual variant interpretation. We found that a minimum of eleven total pathogenic and benign variant controls are required to reach moderate-level evidence in the absence of rigorous statistical analysis.

**Conclusions:** The recommendations and approach to functional evidence evaluation described here should help clarify the clinical variant interpretation process for functional assays. Further, we hope that these recommendations will help develop productive partnerships with basic scientists who have developed functional assays that are useful for interrogating the function of a variety of genes.

## BACKGROUND

The American College of Medical Genetics (ACMG) and the Association for Molecular Pathology (AMP) jointly developed standards and guidelines for assessment of evidence to increase consistency and transparency in clinical variant interpretation [1]. One type of evidence defined in this guideline was the effect of a variant on gene/protein function as determined by a “well-established” functional assay, which provides strong support of a pathogenic or benign impact (rule codes PS3 and BS3, respectively). The full definition is provided in Text Box 1. Functional studies can provide powerful insight into the effect of a variant on protein function and have the capacity to reclassify variants of uncertain significance (VUS) [2], underscoring the need for experimental evidence to be applied accurately and consistently in variant interpretation. However, the ACMG/AMP standards did not provide detailed guidance on how functional evidence should be evaluated, and differences in application of the PS3/BS3 codes represent a major contributor to variant interpretation discordance among clinical laboratories [3].

In response to calls to further standardize variant interpretation [3, 4], the Clinical Genome Resource (ClinGen) established the Sequence Variant Interpretation Working Group (SVI) [5] and condition-specific Variant Curation Expert Panels (VCEPs) to refine ACMG/AMP guidelines for each evidence criterion [6]. To date, six VCEPs have published recommendations, including their assay validation requirements and which assays were ultimately approved for PS3/BS3 evidence application [7–12]. VCEP-approved assays varied greatly and included splicing assays, animal and cellular models, and different in vitro systems [Kanavy et al., submitted]. VCEPs generally approved assays that considered the disease mechanism and most included wild-type controls, but statistical analyses and the inclusion of other controls were less consistent. The VCEPs vary significantly in how they defined which assays were “well-established” [Kanavy et al., submitted], including consideration of parameters such as experimental design, replication, controls, and validation, indicating the subjective nature of assessing the quality and applicability of functional evidence, potentially leading to discordance in variant classification.

In this manuscript, we detail additional guidance developed by the SVI regarding assessment of the clinical validity of functional studies and a provisional framework for the determination of suitable evidence strength levels, with the goal that experimental data cited as evidence in clinical variant interpretation meets a baseline quality level. We expect to further refine these approaches in collaboration with VCEPs as they apply these recommendations moving forward.

## METHODS

In November 2018, during the monthly SVI Working Group conference call, we first outlined our goals of defining what constitutes a well-established functional assay and how functional assay evidence should be structured for computation and curation. In this meeting, we presented a preliminary approach to curating functional evidence and important considerations for assay validation. This process was subsequently presented at the ClinGen Steering Committee in-person meeting in Seattle, WA, in December 2018 for comments and further refinement. The proposed PS3/BS3 evaluation process was then discussed on the SVI Working Group call in March 2019 and again in-person at the American College of Medical Genetics and Genomics (ACMG) meeting in April 2019. Subsequently, a smaller subgroup developed a final version of these recommendations, incorporating feedback from ClinGen biocurators and VCEPs, which were then approved by the SVI Working Group.

We used curated functional evidence from VCEP-developed rule specifications [Kanavy, et al., submitted] and expert opinions throughout the PS3/BS3 criterion refinement process. Feedback from the broader SVI working group, ClinGen Steering Committee, and outside experts were incorporated into the recommendations at multiple stages of development.

To estimate the magnitude of evidence strength that is appropriate for a given assay in the absence of rigorous statistical analysis, we estimated the Odds of Pathogenicity (OddsPath) that could be obtained for a theoretical assay that evaluated various numbers of previously classified controls (see Additional File 1). We treated the proportion of pathogenic variants in the overall modeled data as a prior probability (P_1_), and the proportion of pathogenic variants in the groups with functionally abnormal or functionally normal readouts as posterior probabilities (P_2_). The stringency of the thresholds determining an abnormal versus normal readout is related to the confidence in the assay result. We initially estimated an optimistic OddsPath that could be achieved by a perfect binary classifier, where the readout for all control variants tested is consistent with the variant interpretation (see Supplementary Table 1, Additional File 1). We then sought to estimate a more conservative OddsPath for imperfect assays where one of the control variants had an intermediate or indeterminate readout, but the remaining pathogenic and benign controls would have readouts concordant with their classification (see Supplementary Table 2, Additional File 1) [13, 14]. To circumvent posterior probabilities of zero or infinity, and to account for the possibility that the next variant tested in the assay might have a discordant result, we added exactly one misclassified variant to each set [15]. The OddsPath was estimated for each as: OddsPath=[P_2_ x (1-P_1_)] / [(1-P_2_) x P_1_] [16]. Each OddsPath was then equated with a corresponding level of evidence strength (supporting, moderate, strong, very strong) according to the Bayesian adaptation of the ACMG/AMP variant interpretation guidelines [17].

## POINTS TO CONSIDER AND GENERAL RECOMMENDATIONS

### Physiologic context

The genetic construct and context being evaluated in an assay are important considerations for determining appropriateness for clinical variant interpretation. The assay material being utilized (e.g., patient-derived sample, model organism, cellular *in vivo* or *in vitro* system) should be taken into account when evaluating the validity of a functional assay. When using patient-derived samples, a functional assay evaluates a broader genetic and physiologic background (other variants *in cis* and *in trans*, epigenetic effects, cell type, assay conditions, etc.). For conditions inherited in an autosomal recessive pattern, biallelic variants are required, often in a loss of function mechanism where the penetrance and expressivity of disease manifestations may depend on thresholds of overall protein activity that reflect the cellular/biochemical phenotype arising from a combination of variants and potentially other cellular gene products. In this case, it will be important to distinguish the overall protein activity levels that cause different phenotypes (severe versus mild disease) from the functional assay results that would qualify for variant-level evidence toward a pathogenic or benign interpretation. If a variant is known to be homozygous (either by segregation analysis or exclusion of a large deletion *in trans*), and can be evaluated in multiple unrelated individuals, functional assay evidence from patient-derived material can be interpreted with greater confidence.

> ***Recommendation 1:*** Functional evidence from patient-derived material best reflects the organismal phenotype and, in general, it would be better to use this evidence to satisfy PP4 (specific phenotype) and to delineate the expected disease phenotype in patients with certain combinations of variants or homozygous variants with known pathogenicity. If the curator decides to proceed with evaluating an assay performed in patient-derived material, the level of strength applied should be determined based on validation parameters (see below). In the context of a VCEP, gene-specific guidance should include the required number of unrelated individuals in whom the variant has been tested, in order for the evidence to qualify for variant interpretation.

Typically, model organisms are used to implicate the role of a gene in a disease (e.g., the gene is deleted, interrupted, or an artificial mutation is made to recapitulate a phenotype as evidence of the genetic etiology). Issues related to cost and throughput have typically limited the generation of extensive allelic series intended for the purpose of clinical variant interpretation. In addition, it can be challenging to assess how well the model organism reflects human anatomy/physiology/genetic context, or whether the full phenotype must necessarily be recapitulated in order to satisfy the functional evidence criteria. The genome of the organism may include an orthologous gene (having equivalent or similar function), or the model organism may lack relevant homologs that affect the phenotype in humans, thus affecting the degree to which an artificially introduced genetic variant can cause a relevant phenotype. Even within a given species, measurable phenotypes can vary depending on the genetic background of the organism (e.g., compensatory variation) and therefore studies using more than one strain or line would be preferable, further increasing the cost of such assays. Therefore, the recommendations herein will primarily focus on cellular and biochemical *in vivo* or *in vitro* assays, which are commonly encountered in laboratory evaluations of variants implicated in human disease.

> ***Recommendation 2:*** From the point of view of clinical variant interpretation, evaluation of functional evidence from model organisms should take a nuanced approach, considering the caveats described above. If model organism data are to be used in variant interpretation, the strength of evidence should be adjusted based on the rigor and reproducibility of the overall data provided.

### Molecular consequence

The nature of the variant and the context in which it is studied can significantly affect the assay readout. The effect of the variant on the expressed gene product must be carefully considered when determining the clinical validity of an assay that utilizes an artificially engineered variant. For example, CRISPR-introduced genetic variants in an otherwise normal genomic context will use the endogenous cellular transcriptional and splicing machinery, although off-target effects must be carefully considered. In contrast, transient expression of cDNA constructs, which usually contain artificial promoters and other regulatory sequences that can result in variant overexpression, should be carefully standardized using controls to ensure that the overexpression does not mask the true effects of variants. Nonsense and frameshift variants that result in premature termination codons before the 3’-most 50 nucleotides of the penultimate exon are expected to undergo nonsense-mediated decay (NMD) and eliminate the mRNAs [18, 19]; therefore, studying such variants in the context of cDNA or systems where NMD is not active may not reflect the endogenous situation. Similarly, the effects of a nucleotide substitution or other in-frame variant on splicing cannot be assessed using a cDNA construct. On the other hand, when the variant results in an expressed protein with an in-frame deletion or a single nucleotide substitution, an engineered cDNA construct may reasonably reflect the functional impact, at least at the protein level.

> ***Recommendation 3:*** While testing variants in a more natural genomic context is preferable, it is not a requirement of a well-validated assay. Instead, one should consider how the approach impacts the interpretation of the results and take into account whether the study controls for these limitations when assigning strength of evidence.

Since an individual functional assay may not fully capture all gene or protein functions relevant to disease pathogenesis, a “normal” result in a laboratory assay may simply reflect that the functional effect of the specific variant was not suitably assayed in the experiment. Therefore, in order to determine when, and at what strength, to apply the BS3 criterion, it is essential to understand how well the assay captures the molecular consequence of the variant and its impact on the expressed protein or functional domain. A more complete assessment of protein function permits scoring the result as having a benign effect, whereas an assay that is limited to a specific domain or functional readout may provide less strong evidence for having a benign effect. It should also be noted that a missense or synonymous variant that does not affect protein function might still have a negative impact by introducing a cryptic splice site [20]. These caveats should be taken into account when deciding whether to apply BS3, and at what strength.

Messenger RNA splicing is a complex process, and clinical variant interpretation can take into account both predictive and laboratory evidence. RNA splicing assays, developed using the endogenous genomic context, or using artificial mini-gene assays, can be useful to determine the impact of variants on splicing integrity. However, unlike protein assays, the readout (e.g., exon skipping, or intron retention) does not necessarily correlate with protein function. For example, abnormal splicing of the last exon might lead to a truncated protein whose function is still intact. In general, abnormal splicing can have heterogenous outcomes with respect to mRNA fate and the protein-reading frame. Abnormally spliced transcripts might undergo NMD, while other abnormal transcripts can lead to a shortened or truncated protein with or without functional consequences [21]. The relative transcript abundance of various splice isoforms in different cell types may also affect the downstream pathophysiological impact.

Because RNA splicing assays do not provide a direct measure of protein function, additional recommendations are needed to determine the applicability of splicing assays to satisfy PS3/BS3 versus PVS1 (loss-of-function). For canonical ±1,2 splice variants, PVS1 application is based on the predicted impact of a variant on mRNA stability and protein-reading frame whereas a functional assay may conclusively demonstrate abnormal splicing and confirm a loss of function impact. Additional data and considerations are needed to determine the appropriate aggregate strength of PVS1 and PS3 in the scenario that functional data is present and supports PVS1 application. Similarly, splicing assays could be used to bolster support for *in silico* predictions for variants outside the canonical ±1,2 splice sites. An SVI subgroup is currently working on recommendations for the incorporation of predictive and functional evidence of altered splicing into the ACMG/AMP framework. For variants impacting protein length that are not predicted to lead to loss-of-function, such as in-frame exon skipping due to abnormal splicing or a large in-frame deletion, the change in protein length alone could be used to justify application of PM4, while application of PS3/BS3 could also be appropriate if a functional assay examined the protein function of the resulting product.

### Terminology

Standardized, structured language can improve communication and transparency across clinical laboratories, physicians, and patients. Uniform terminology should be used to describe the readout of a laboratory assay of protein function and document the curation of functional evidence. As such, the variant-level results of functional assays should not be categorized as “pathogenic” or “benign,” since these falsely equate functional impact with a clinical determination that involves a number of other evidence lines. In addition, terms describing assay results as “deleterious”, or “damaging” can be confusing since their meanings are greatly context-dependent and generally only apply when loss of function is the mechanism of disease. For example, in conditions where the mechanism involves gain of function, a variant may be damaging or deleterious to the organism but not to protein activity as measured in a functional assay. Establishing standardized language to describe assay readout is an important step to prevent the misinterpretation of published data and to reduce inter-laboratory discordance with respect to PS3/BS3 application [3, 22].

> ***Recommendation 4:*** The terms “functionally normal” or “functionally abnormal” should be used to describe the functional impact of a variant as measured in a given assay. Further granular specifications should be used to describe the “functionally abnormal” impact (i.e., complete loss of function, partial loss of function/intermediate effect/hypomorphic, gain of function, dominant-negative) as outlined by Spurdle, et al. [22]. The final assessment of the evidence should take into account both the functional effect in the assay and the mechanism of disease (see below).

### CLIA laboratory developed tests

The 2015 ACMG/AMP guidelines assert that “functional studies that have been validated and shown to be reproducible and robust in a clinical diagnostic laboratory setting are considered to be the most well established” [1]. All tests conducted in a Clinical Laboratory Improvement Amendments (CLIA) laboratory or with a commercially available kit are subject to analytical validation for in-house use. However, these assays should also be evaluated for the strength of evidence based on the controls used, as detailed below. One should also consider that *in vitro* assays developed in CLIA laboratories that are conducted with patient samples for diagnostic use [23] may not necessarily provide variant-level evidence relevant to interpretation (see Recommendation 1, above). Data from research laboratories are not subjected to specific regulatory oversight and thus may be validated to different degrees, though any *in vivo* or *in vitro* study can satisfy PS3/BS3 criteria with a strong level of evidence if it demonstrates the appropriate validation.

> ***Recommendation 5:*** The entity performing a functional assay should not govern whether PS3/BS3 criteria are satisfied, or at what strength. This determination should be based primarily on the validation of the assay, including the use of appropriate laboratory controls as well as clinical validation controls (as described below).

### Experimental controls and clinical validation controls

Good laboratory practice is essential for application of functional evidence in clinical variant interpretation. Every experiment should include internal controls that demonstrate the dynamic range of the assay (e.g., the readout of the assay with wild type and null effect). In some cases, the readout may be normalized to a wild type value, which should generally be run in the same conditions as the variants being tested to avoid a batch effect. Well-conducted experiments typically use technical replicates that control for the random differences associated with protocol or instrument-associated variation, to demonstrate reproducibility of the result within a given experiment. Similarly, biological replicates (e.g., different colonies, cells, aliquots, or animals) are included to control for random biological variation in parallel measures of unique biological samples and to demonstrate reproducibility of the result between instances of the same experiment. Biological replicates are more important for understanding the variance within a population, while technical replicates can reduce measurement error [24].

Furthermore, well-validated assays are benchmarked by including known pathogenic and known benign variants that establish the ranges of assay readout for these classes of variants and define the thresholds beyond which the result can be considered functionally abnormal, indeterminate, or functionally normal. It is important to note that the clinical interpretation of these validation control variants should reach a pathogenic/likely pathogenic or benign/likely benign interpretation using lines of evidence independent of functional data, so as to avoid circularity in defining the assay’s predictive value. The number of controls required depends on the dynamic range of the assay and the variance of each replicate; controls should also be relevant to the disease mechanism (such as gain-of-function or loss-of-function) and the type of variant under consideration (e.g., missense controls for evaluating missense variants of uncertain significance). For genes associated with multiple disorders through different mechanisms, an assay validated for one disorder may not necessarily be applied universally to analyze variant effect in other disorders if the mechanisms of disease are different. Variants in the Genome Aggregation Database (gnomAD)[25] that have population allele frequencies exceeding the threshold for BA1 or BS1, but have not yet been added to the Clinical Variant Database (ClinVar), could serve as a source of benign controls. Additionally, one could consider if pathogenic or benign controls from different genes related via a disease mechanism and functional pathway could be used at a lesser strength of evidence.

Many previously published assays do not identify known benign or known pathogenic variant controls, or may have only tested a few variant controls in the same assay. To address this, it may be possible for analysts to assemble these controls from multiple specific instances of the same general class of assay. Any tested variant that could be classified as likely benign/benign (LB/B) or likely pathogenic/pathogenic (LP/P) without functional criteria would qualify as a control for the determination of evidence strength. The assay readout for each of these variants, as tested across multiple instances of the same general class of assay, can be plotted together in order to set thresholds for normal, intermediate, and abnormal function (Figure 1).

**Figure 1:**
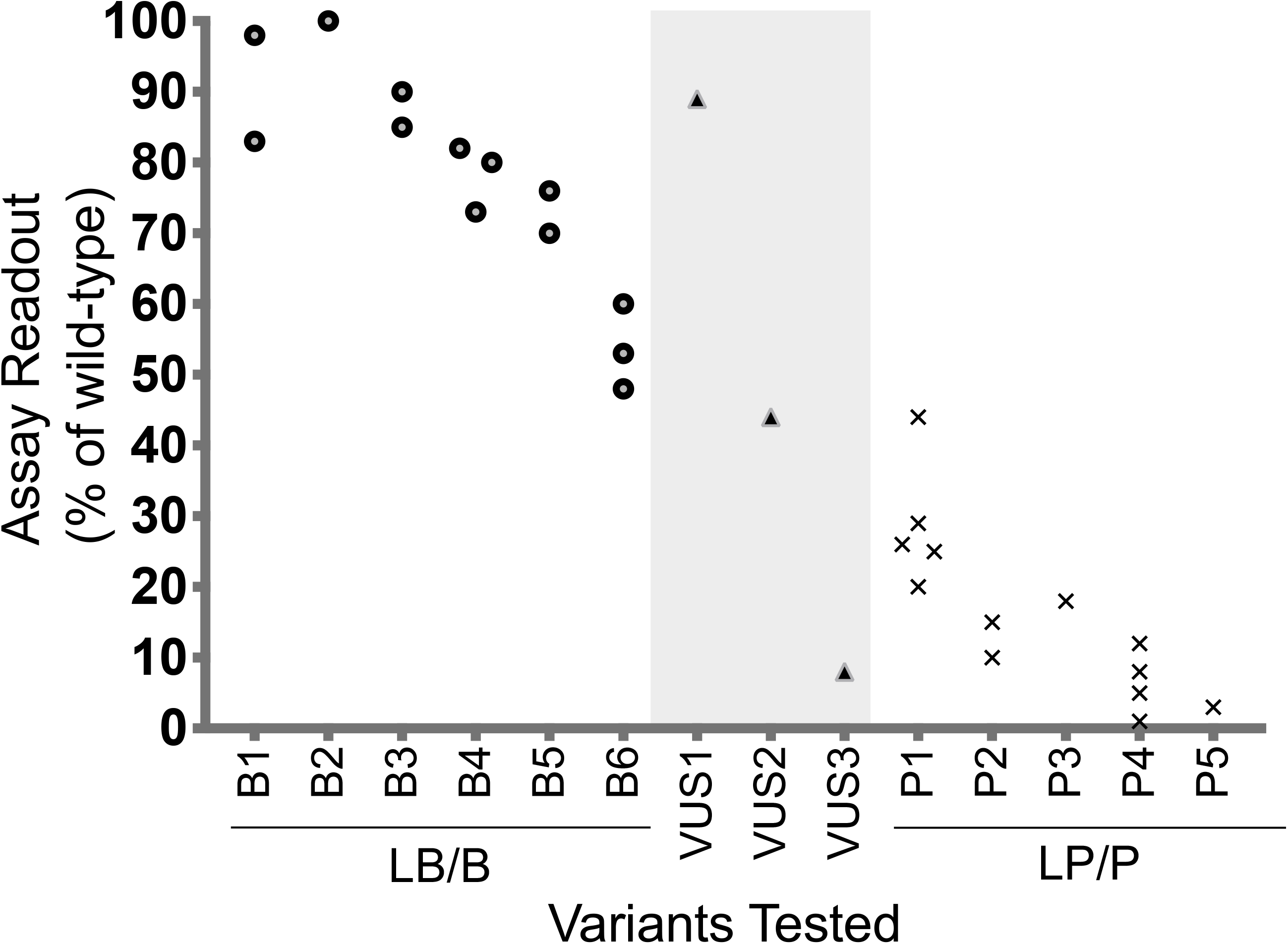
Assembly of variant controls to set readout thresholds for normal and abnormal function. Readout values across multiple specific instances of the same type can be plotted for any tested variant that reaches a likely benign/benign (LB/B) or likely pathogenic/pathogenic (LP/P) classification without PS3 or BS3 criteria. Each point on the plot represents the assay readout from a specific instance of an assay for the variant listed on the x-axis. Multiple points for the same variant indicate that the variant was tested in multiple specific instances of the same general class of assay. In this example, all LB/B variant controls (B1-B6) had readouts above 60%, with the exception of variant B6. When setting a readout threshold above which the readout is considered normal function, curators may draw this threshold at 60% and consider B6 to have an indeterminate readout. All LP/P variant controls (P1-P5) had readouts below 30%, with the exception of one specific instance for variant P1. With just 1 LB/B control variant with an indeterminate readout from a total of 11 variant controls (6 LB/B and 5 LP/P), PS3_moderate can be applied to variants with a readout indicating abnormal function and BS3_moderate can be applied to variants with a readout indicating normal function (see Supplemental Table 2, Additional File 1). Variants of uncertain significance (VUS) tested on the same class of assay are plotted in the middle of the graph (indicated by light gray shading). VUS1 has an assay readout in the range of LB/B controls and would be above the threshold for normal function, so BS3_moderate could be applied. VUS3 has an assay readout consistent with LP/P control variants, below the threshold for abnormal function, so PS3_moderate could be applied. VUS2 has an indeterminate assay readout, so neither PS3_moderate nor BS3_moderate can be applied for this variant.

## PROVISIONAL FRAMEWORK FOR FUNCTIONAL EVIDENCE EVALUATION AND APPLICATION

The SVI Working Group recommends that evaluators use a four-step process to determine the applicability and strength of evidence of functional assays for use in clinical variant interpretation: 1. Define the disease mechanism; 2. Evaluate applicability of general classes of assays used in the field; 3. Evaluate validity of specific instances of assays; 4. Apply evidence to individual variant interpretation. Unlike the ACMG/AMP guidelines [1], in which well-established functional studies can provide a default “strong” level of evidence (PS3/BS3), the SVI recommends that evaluation of functional assays should start from the assumption of no evidence, and that increasing clinical validation can allow application of evidence in favor of a pathogenic or benign interpretation at a level of strength (supporting, moderate, strong) concomitant with the demonstrated validation metrics as described below.

### 1. Define the disease mechanism

In order for functional assays to be useful in clinical variant interpretation, the underlying gene-disease mechanism must be reasonably well understood. The VCEP or individual interpreting variants in a given gene should first delineate this mechanism to determine what functional assays can be considered applicable. This is an important first step since some genes are associated with different diseases depending on the mechanism (e.g., gain-of-function vs. loss-of-function). A structured narrative using ontologies or other specific terms can be used to describe the gene-disease mechanism (Text Box 2).

### 2. Evaluate applicability of general classes of assays used in the field

Next, the general types or classes of assays used in the field should be defined and documented, including the model system, experimental method, and functional outcome being evaluated. The defined gene-disease mechanism should guide an evaluation of how well a general class of assay models pathogenesis (e.g., loss of function, gain of function, specific pathway output, etc.). Relative strengths and weaknesses of the model system should be assessed and disease-specific assertions regarding the appropriateness of animal, cellular, and *in vitro* models should be addressed (see above: physiologic context, molecular consequence). The purpose of this step is to delineate the types of assays that are deemed appropriate (if sufficiently validated) for use in clinical variant interpretation. It is important to reiterate that the strength of evidence is not determined by the class of assay but rather by the validation metrics (specified in step three).

For expert groups that are establishing gene-specific guidance, we also recommend that they refrain from making blanket statements limiting the general classes of assay that are deemed valid or applicable, and should not cap the strength of evidence based on the class of assay. In some cases, a VCEP may wish to endorse a particular type of assay that could be used for variant interpretation if developed in the future.

### 3. Evaluate validity of specific instances of assays

For the general classes of assay that are deemed applicable, the curator should next evaluate specific instances of those assays as performed by various groups. Many different laboratories may generate functional evidence using the same general class of assay, but given the differences in the specific methods used and the level of validation provided by each group, evaluation of each individual assay iteration is required before data can be applied in a clinical interpretation (see above section on experimental controls and clinical validation controls). Assays having sufficient numbers of validation controls to calculate positive predictive value or determine the OddsPath provide the most robust functional assay evidence [17]. Without this level of clinical validation, the predictive value of the assay is limited. A provisional framework for this evaluation is shown in Figure 2.

- Functional evidence should not be applied in the following scenarios unless the dynamic range of the assay and the thresholds for defining a functionally normal, indeterminate, or functionally abnormal result are extremely well understood:

- Assays that do not include both negative (normal or wild-type) and positive (abnormal or null) controls.
- Assays that do not include technical and/or biological replicates.
- Supporting level evidence in favor of pathogenicity (PS3_supporting) or benign interpretation (BS3_supporting) may be applied in the following scenarios:

- Assays that include experimental controls and replicates but have 10 or fewer validation controls to assess the ability of the assay readout to distinguish pathogenic from benign variants (see Supplementary Table 2, Additional File 1).
- Classes of assays that have been broadly accepted historically, previously validated, or provided as a kit with defined performance characteristics, but where controls and replicates are not documented for the specific instance of the assay.
- Moderate level evidence in favor of pathogenicity (PS3_moderate) or benign interpretation (BS3_moderate) may be applied in the following scenarios:

Assays with at least 11 total validation controls including a mix of benign and pathogenic variants, but no formal statistical analysis of the ability to discriminate between pathogenic and benign variants (see Supplementary Table 2, Additional File 1)
- Any level of evidence in favor of pathogenicity may be applied when rigorous statistical analysis enables a formal OddsPath to be calculated, with the strength of evidence correlating to the calculated OddsPath (Table 1).
- Evidence in favor of a benign interpretation up to a strong level (BS3) may be applied when rigorous statistical analysis enables a formal OddsPath to be calculated, with the strength of evidence correlating to the calculated OddsPath (Table 1).

**Figure 2:**
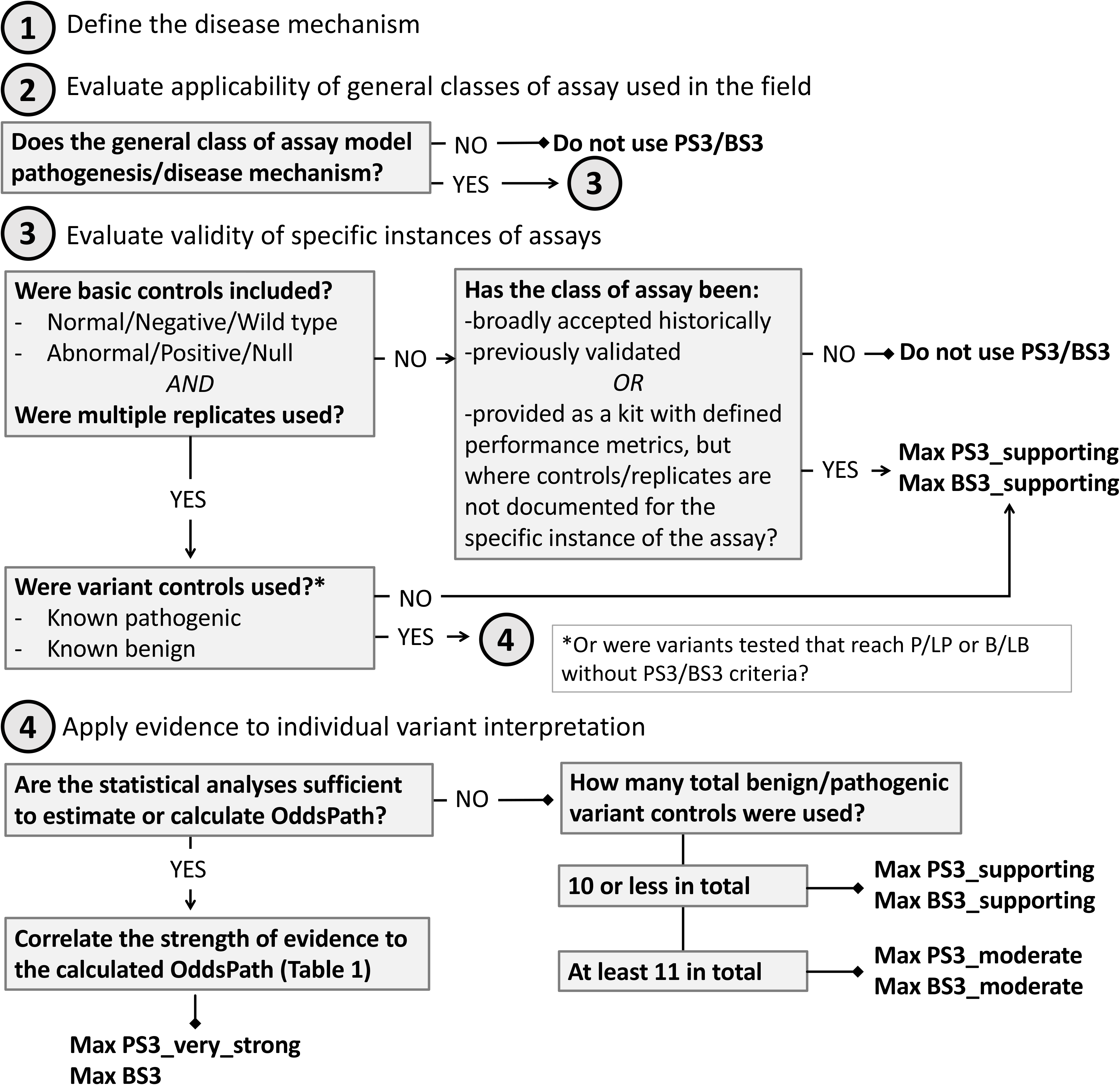
Decision tree for evaluation of functional data for clinical variant interpretation. The SVI Working Group recommends that evaluators use a four-step process to determine the applicability and strength of evidence of functional assays for use in clinical variant interpretation (evidence codes PS3/BS3): 1. Define the disease mechanism; 2. Evaluate applicability of general classes of assay used in the field; 3. Evaluate validity of specific instances of assays; 4. Apply evidence to individual variant interpretation.

**Table 1:** Evidence strength equivalent of Odds of Pathogenicity.

| Odds of Pathogenicity (OddsPath) | Evidence strength equivalent |
| --- | --- |
| <0.053 | BS3 |
| <0.23 | BS3_moderate* |
| <0.48 | BS3_supporting |
| 0.48-2.1 | Indeterminate |
| >2.1 | PS3_supporting |
| >4.3 | PS3_moderate |
| >18.7 | PS3 |
| >350 | PS3_very_strong |

\*Since there are no moderate strength benign evidence codes included in the Richards, et al. guidance, moderate level evidence is equivalent to two instances of supporting-level benign evidence [17]

VCEPs should document the specific assay instances that qualify (and why) and the specific instances of assays that do not qualify (and why). Documentation should include PMID or other universal reference to the source of the assay evaluated (e.g., DOI), the type of assay readout (qualitative/quantitative) and units, the range of assay results that qualify for a given strength of evidence according to level of validation as above, and the range in which the assay result is indeterminate with respect to PS3/BS3.

### 4. Apply evidence to individual variant interpretation

Once the specific instance of an assay has been evaluated as a whole, the results from that assay for a given variant can be applied as evidence in variant interpretation.

- If the assay demonstrates a functionally abnormal result consistent with the mechanism of disease, the PS3 criterion can be applied at a level of strength based on the degree of validation detailed above.
- If the assay demonstrates a functionally normal result, the BS3 criterion can be applied at a level of strength based on the degree of validation detailed above.
- Variants demonstrating an intermediate level of impact on function merit special consideration, as this could be because the assay does not fully reflect the protein function (decreasing strength applied to the assertion), or may provide evidence supporting a hypomorphic or partial loss-of-function effect, such as in a condition with incomplete penetrance and/or less severe expressivity. Consideration of disease mechanism should help guide the appropriate level of strength to be applied for these types of variants.

When PS3/BS3 are applied by any variant analyst, documentation of the supporting evidence should reference the strength of clinical validation of the functional assay.

#### Stacking Evidence

When multiple functional assay results are available for a single variant (different instances of the same class of assay performed by different laboratories, or multiple lines of evidence from different classes of assay), the evaluator should apply evidence from the assay that is the most well-validated and best measures the disease mechanism.

- For a variant analyzed by multiple assays (belonging to the same or different class):

- If the results are consistent (both show a functionally abnormal effect, or both show a functionally normal effect), apply PS3/BS3 at the level of strength appropriate for the most well-validated assay.
- If the results are conflicting, the assay that most closely reflects the disease mechanism and is more well-validated can override the conflicting result of the other and evidence should be applied at the strength indicated by the assay’s validation parameters. If the assays are essentially at the same level of validation, conflicting functional evidence should not be used in the interpretation of the variant.
- The committee did not reach consensus on whether results from different classes of functional assay could be combined (e.g. applying two pieces of supporting level evidence from different assay classes to reach PS3_moderate). The primary concern with this approach is that it is extremely difficult to ascertain that two assays are measuring independent functions and that this would lead to double counting the same evidence regarding variant function. Another concern is that stacking evidence from multiple assays could lead to a conflated interpretation of the disease risk for a particular variant (e.g. two PS3_supporting could be interpreted as concordant evidence that the variant confers moderate disease risk; alternatively, two PS3_supporting results could stack to PS3_moderate as a high-risk variant). On the other hand, if the assays are measuring different functions, the evidence may be complementary and increase confidence in the overall result, especially for the assertion of BS3 criteria. Variant curators and expert groups will need to decide how to best proceed, keeping in mind the cautions regarding the double counting of evidence.

## NEXT STEPS AND FUTURE CONSIDERATIONS

### Framework evolution

This provisional framework for the evaluation and application of functional evidence in clinical variant interpretation represents the first important steps towards reducing discordance in the use of PS3/BS3 criteria. Moving forward, this approach will be tested with a range of diverse disorders in collaboration with ClinGen VCEPs. We recognize that many historical publications may not meet the specifications outlined here, which will limit our ability to apply these assays as strong evidence in the ACMG/AMP variant interpretation framework, though they may still qualify for supporting-level evidence if performed rigorously and with appropriate laboratory controls. Undoubtedly, many other kinds of evidence will be re-weighted as the ACMG/AMP guidelines are revised and this provisional framework will evolve alongside these updates.

### Bayesian adaptation

As the field moves to develop assays with sufficient controls and validation to permit the calculation of an OddsPath, more quantitative approaches for stacking evidence and assigning evidence strength may be adopted, as outlined in the Bayesian adaptation of the ACMG/AMP variant interpretation framework [17]. This quantitative method will reconcile conflicting benign and pathogenic evidence, which is common when considering the results of multiple functional assays, and will help reduce the number of VUS. Furthermore, many assays provide continuous quantitative measures of protein function, and converting their numeric readout to a binary PS3/BS3 interpretation can obscure the richness of those data. Using a more quantitative Bayesian system could convert raw data to OddsPath that more completely capture the assay results. This would be especially useful for hypomorphic variants that have an intermediate effect on normal protein function.

### Multiplexed functional assays

While typical functional assays cited as evidence in variant curations analyze relatively few variants [Kanavy, et al., submitted], new multiplexed assays can analyze thousands of variants in a single experiment [26–28]. This kind of increased throughput facilitates the reproducibility, replication, and assay calibration using many definitive pathogenic and benign variant controls. These metrics are required to determine assay sensitivity and specificity, which can then guide the interpretation of assay readout according to thresholds set by known benign and known pathogenic variant performance. Similarly, thresholds could be drawn based on OddsPath to apply different strengths of evidence based on the specific assay result. Multiplexed assays are still heavily dependent upon the existence of well-characterized pathogenic and benign variants for assay validation. The availability of allelic variant controls may be limited for some genes, but threshold determination may still be feasible depending on the assay’s dynamic range and the distribution of results relative to null and wild-type controls (including variants with high allele frequency incompatible with a pathogenic role for rare Mendelian diseases). In the future, these large datasets of functional evidence could be ingested into the Variant Curation Interface (VCI) or Evidence Repository and made available to variant curators in an automated fashion alongside pre-determined thresholds for interpretation and strength assignment, expediting the curation process. Such an automated repository could re-assess sensitivity and specificity automatically as more variants are added. It is important to note that even if the functional data reach an OddsPath equivalent to very strong evidence, the functional evidence criteria are not stand-alone evidence for either a benign or pathogenic classification and at least one other evidence type (e.g., PS4, prevalence in affected individuals is significantly increased relative to controls) is required to reach a pathogenic classification.

### Prioritization methods for functional assay development and validation

As it is time-consuming and expensive to develop and sufficiently validate novel functional assays, effort and resources should be directed to have the greatest clinical benefit. One could prioritize assays that would examine genes with the greatest number of genetic tests performed or individuals tested annually, or focus on the genes with the greatest proportion of VUS that could be adjudicated with functional evidence [29]. Alternatively, one could focus on genes associated with highly actionable conditions, where a change in variant interpretation might dramatically change medical management (e.g., *BRCA2* VUS would be reclassified as likely pathogenic with functional evidence, leading to increased early surveillance and recommendations regarding cancer prophylaxis and management).

We hope that these recommendations will help develop productive partnerships with basic scientists who have developed functional assays that are useful for interrogating the function of a variety of different genes [30]. Realistically, many researchers may not envision a use for their assays in clinical variant interpretation, and may not recognize the need for extensive validation when applying this evidence clinically (nor possess the expertise to independently determine the clinical interpretation of variants in the gene of interest). We look forward to partnerships between VCEPS and basic scientists to apply the results of *in vitro* and *in vivo* tests in clinical variant interpretation. Publishing and/or submitting these results to ClinGen along with appropriate documentation of validation and thresholds for interpretation will greatly enhance curation and application of these data. Greater awareness of the validation requirements, especially the use of an allelic series containing known pathogenic and known benign variants to evaluate the predictive value of the assay, may enable such assays to be used for clinical interpretation more broadly in the future.

## LIST OF ABBREVIATIONS

ACMG: American College of Medical Genetics and Genomics
AMP: Association for Molecular Pathology
B: benign
BA1: allele frequency data as stand-alone evidence of benign impact
BS1: allele frequency greater than expected for disease, strong evidence of benign impact
BS3: well-established functional studies provide strong support of a benign effect
cDNA: complementary deoxyribonucleic acid
CLIA: Clinical Laboratory Improvement Amendments
ClinGen: Clinical Genome Resource
ClinVar: Clinical Variant Database
CRISPR: clustered regularly interspersed short palindromic repeats
DOI: Digital Object Identifier
gnomAD: Genome Aggregation Database
LB: likely benign
LP: likely pathogenic
mRNA: messenger ribonucleic acid
NMD: nonsense-mediated decay
OddsPath: odds of pathogenicity
P: pathogenic
PM4: protein length changes as a result of in-frame deletions/insertions in a nonrepeat region or stop-loss variant, moderate-level evidence of pathogenic impact
PMID: PubMed Identifier
PP3: computational, supporting-level evidence of pathogenic impact
PP4: phenotype is highly specific for disease, supporting-level evidence for pathogenicity
PS3: well-established functional studies providing strong support of a pathogenic effect
PS4: prevalence in affected individuals is significantly increased relative to controls, strong evidence of pathogenic impact
PVS1: null variant where loss-of-function is a known mechanism of disease, very strong evidence of pathogenicity
RT-PCR: real-time polymerase chain reaction
SVI: Sequence Variant Interpretation Working Group
VCEP: Variant Curation Expert Panel
VCI: Variant Curation Interface
VUS: variant of uncertain significance

## Declarations

### Ethics approval and consent to participate

Not applicable.

### Consent for publication

Not applicable.

### Availability of data and material

Not applicable.

### Authors’ contributions

SEB and JSB conceived the study. SVT developed the model and analysis of evidence strength. SEB, JSB, AAT, SMH, SMM, SVT and DMK drafted the manuscript. SEB, JSB, SMM, AAT, SMH, AAT, LMS, CDH, MSG, GRC, FJC, XL, and LGB reviewed and edited the manuscript draft. All authors read and approved the final manuscript.

## Acknowledgements

ClinGen Sequence Variant Interpretation Working Group Members: Ahmad Abou Tayoun, Al Jalila Children’s Specialty Hospital, Dubai, UAE; Jonathan S. Berg, University of North Carolina, Chapel Hill, NC; Leslie G. Biesecker, co-chair, National Human Genome Research Institute, National Institutes of Health, Bethesda, MD; Steven E. Brenner, University of California, Berkeley, Berkeley, CA; Garry R. Cutting, Johns Hopkins University School of Medicine, Baltimore, MD; Sian Ellard, University of Exeter Medical School, Exeter, UK; Marc S. Greenblatt, University of Vermont, Larner College of Medicine, Burlington, VT; Steven M. Harrison, co-chair, Broad Institute of MIT/Harvard, Cambridge, MA; Izabela Karbassi, Quest Diagnostics, Athena Diagnostics, Marlborough, MA; Rachel Karchin, Johns Hopkins University, Baltimore, MD; Jessica L. Mester, GeneDx, Inc., Gaithersburg, MD; Anne O’Donnell-Luria, Boston Children’s Hospital, Boston, MA; Tina Pesaran, Ambry Genetics, Aliso Viejo, CA; Sharon E. Plon, Baylor College of Medicine, Houston, TX; Heidi Rehm, Massachusetts General Hospital, Boston, MA; Sean Tavtigian, University of Utah School of Medicine, Salt Lake City, UT; Scott Topper, Color Genomics, Burlingame, CA.

## Authors’ information

The opinions and recommendations expressed here are those of the authors and do not necessarily reflect the policies of any institutions or other organizations to which they are affiliated.

**Text Box 1: Text of Original ACMG/AMP Recommendation for Functional Assays, Reproduced with Permission** [1] “Functional studies can be a powerful tool in support of pathogenicity; however, not all functional studies are effective in predicting an impact on a gene or protein function. For example, certain enzymatic assays offer well-established approaches to assess the impact of a missense variant on enzymatic function in a metabolic pathway (e.g., α-galactosidase enzyme function). On the other hand, some functional assays may be less consistent predictors of the effect of variants on protein function. To assess the validity of a functional assay, one must consider how closely the functional assay reflects the biological environment. For example, assaying enzymatic function directly from biopsied tissue from the patient or an animal model provides stronger evidence than expressing the protein *in vitro*. Likewise, evidence is stronger if the assay reflects the full biological function of the protein (e.g., substrate breakdown by an enzyme) compared with only one component of function (e.g., adenosine triphosphate hydrolysis for a protein with additional binding properties). Validation, reproducibility, and robustness data that assess the analytical performance of the assay and account for specimen integrity, which can be affected by the method and time of acquisition, as well as storage and transport, are important factors to consider. These factors are mitigated in the case of an assay in a Clinical Laboratory Improvement Amendments laboratory–developed test or commercially available kit. Assays that assess the impact of variants at the messenger RNA level can be highly informative when evaluating the effects of variants at splice junctions and within coding sequences and untranslated regions, as well as deeper intronic regions (e.g., messenger RNA stability, processing, or translation). Technical approaches include direct analysis of RNA and/or complementary DNA derivatives and *in vitro* minigene splicing assays.”

**Text Box 2:** Components of the structured narrative describing the gene-disease mechanism.

- Gene name: HUGO Gene Nomenclature Committee (HGNC) gene symbols [31]
- Associated disease: Monarch Disease Ontology (MONDO) terms [32]
- Mode of inheritance: structured MONDO terms

- Autosomal dominant (HP:0000006)
- Autosomal recessive (HP:0000007)
- Mitochondrial (HP:0001427)
- X-linked (HP:0001417)
- Undetermined (HP:0000005)
- Molecular mechanism of disease pathogenesis:

- Loss of function
- Gain of function
- Dominant negative
- Biological pathways: Gene Ontology (GO) terms [33, 34]

## Additional Files

File name: Additional File 1.xlsx

File format: Excel (.xlsx)

Title of data: Odds of Pathogenicity (OddsPath) by performance of classified variant controls Description of data: Additional File 1 contains two tables modeling the Odds of Pathogenicity and strength of evidence based on the number of known benign and known pathogenic variants tested and the ability of the assay to correctly classify those variants, either as a perfect binary (normal/abnormal) readout (Supplementary Table 1) or permitting one variant control (either benign or pathogenic) to have an indeterminate readout (Supplementary Table 2).

